# Mathematical model of a cell membrane

**DOI:** 10.1101/2023.11.27.568933

**Authors:** Nick Gorkavyi

## Abstract

Bimolecular cell membranes play a crucial role in many biological processes and possess a unique set of physical properties. Bimolecular membranes and monomolecular films can be considered as a “two-dimensional fluid” because the diffusion of molecules along the membrane or film is a hydrodynamic process. On the other hand, the bending of the cell membrane is controlled by its stiffness and elastic tension. The aim of this work is to adapt the Navier-Stokes hydrodynamic equations, obtained using the classical Chapman-Enskog method, to the case of two-dimensional membranes. The hydrodynamic equation system is complemented by an elasticity equation for the bending oscillations of the membrane. The obtained system of equations for the dynamics of the cell membrane is linearized for the case of disturbances with small amplitude. Dispersion equations for stable and unstable linear oscillations of cell membranes are investigated, and conditions for the onset of instabilities are derived.

## 1. Introduction

Biological systems in which hydrodynamic diffusion of molecules is often accompanied by chemical reactions tend to exhibit collective instabilities and the formation of spatially regular structures. Well-known examples include the Belousov-Zhabotinsky reaction and Turing instability (Murray, 1977; Belintsev, 1991). Collective processes and instabilities develop in organisms at various levels, ranging from the dynamics of molecules and ions on cell membranes to morphogenetic instabilities shaping the organs of animals and determining the heterogeneous coloring of skins (Murray, 1977). Experimental data demonstrate spatial structures on the cell membrane, such as metal-ion-induced lipid phase separation and “purple patches” of bacteriorhodopsin on the cell membranes of *Halobacterium halobium* (see, for example, Garret and Grisham, 1995a, 1995b).

Cell membranes, consisting of a double layer of lipids (see, for example, Chernomordik and Kozlov, 2008), possess unique properties Along the membrane surface, lipid and protein molecules move quite freely, akin to a fluid - hence, the membrane is sometimes referred to as a “two-dimensional fluid.” However, the bending of the cell membrane is controlled by its stiffness and elastic tension, and therefore, it is described by an elasticity equation (Belintsev, 1991).

Various theoretical descriptions of the cell membrane and other biological environments using hydrodynamic or diffusion equations, as well as elasticity equations, are detailed in numerous articles and monographs. Different scientific groups are developing analytical and numerical models of the cell and its membrane based on kinetic theory, diffusion equations, thermodynamics and electrodynamics (Helfrich, 1973; Evans and Yeung, 1994; Rochal et al., 2005; Kruse et al., 2005; Liang and Mahadevan, 2009; Reigada et al., 2010; Pinkwart et al., 2019; Marrink et al., 2019) (see also references in these works). The equations in these models, as well as the methods of their derivation, significantly differ from each other due to the diverse objectives set by the authors and the complexity of dynamic processes in biomembranes. The developed models are often incomplete — for example, based on diffusion hydrodynamic equations but lacking an equation for temperature. The diversity of models for cell membranes and the methods used to derive them complicates the comparison of results obtained by different authors.

The Navier-Stokes hydrodynamics equations with phenomenological transport coefficients (viscosity and thermal conductivity) are well studied and experimentally confirmed (Landau and Lifshitz, 1987). Chapman and Enskog developed a method to derive hydrodynamics equations from the Boltzmann kinetic equation, yielding analytical expressions for viscosity and thermal conductivity coefficients (Chapman and Cowling, 1991; Hirschfelder et al., 1954). This method was later generalized for plasmas and ionized gases with chemical reactions (Braginsky, 1963; Ferziger and Kaper, 1972). The Navier-Stokes equations have also been successfully applied to granular media.

It is of interest to obtain a complete system of differential hydrodynamic equations to describe transport processes in cell membranes. Such a system, derived using universally accepted methods, would serve as a reliable starting point for comparing various models. The hydrodynamic equation system for cell membranes needs to be supplemented with an equation describing its bending. Mathematical models for the oscillations of thin plates have long been developed and firmly established in elasticity theory (Landau et al., 1986). Therefore, within the discussed approach, an analogous equation for the bending oscillations of the cell membrane can be formulated. The combination of a unified system of hydrodynamic equations for processes in the plane of the real cell membrane and an elasticity equation for its bending has been previously considered (Helfrich, 1973; Belintsev, 1991).

A comprehensive system of equations for the dynamics of cell membranes will enable the exploration of diverse oscillations and instabilities in the membrane, as well as identify promising directions for constructing more complex models.

Solving the hydrodynamic equation system augmented with the elasticity equation for membrane oscillations is extremely challenging—both analytically and numerically. However, for the simple case of linear oscillations, a dispersion equation can be derived, allowing the investigation of stable and unstable membrane oscillations. The dispersion equation method is widely used in biology (Murray, 1977; Belintsev, 1991), plasma physics and astrophysics (Fridman and Gorkavyi, 1999). This method is applied to analyze oscillations in predator-prey populations and to study instabilities in systems of biochemical reactions with diffusion (Murray, 1977).

The objectives of this work include obtaining and studying:

a. Navier-Stokes-type hydrodynamic equations describing cell membranes and monomolecular films as a two-dimensional fluid;
b. Elasticity equation describing transverse oscillations of a homogeneous cell membrane as a plate with stiffness and elasticity;
c. Dispersion equation for linear oscillations of membranes and films;
d. Specific branches of oscillations—both stable and unstable ones. Deriving criteria for some instabilities of the membrane, including those associated with malaria and several viral infections.

## 2. Mathematical model

Let’s consider the cell membrane as a liquid system in the (X, Y) plane and an elastic plate in the case of Z-bending. The most general equations for hydrodynamics and elasticity can be taken from published monographs and papers (Landau and Lifshitz, 1987; Landau et al., 1986; Gorkavyi et al., 1986). The development of transport equations for a thin layer is detailed in the book (Fridman and Gorkavyi, 1999). The key distinction lies in using two-dimensional hydrodynamics for the cell membrane, where the hydrodynamic velocity and temperature lack Z-components.

It can be demonstrated that two-dimensional hydrodynamic equations, in contrast to three-dimensional hydrodynamic equations (Landau and Lifshitz, 1987), have different coefficients in the energy equation (3/2 is replaced by 1 for 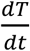) and in the viscosity stress tensor (3/2 is replaced by 1 for the Kronecker delta symbol *δ*_*ik*_). Without these replacements, the discussed system of equations would be applicable not to monomolecular or bimolecular membranes but to macromolecular thin layers, for example, epithelia.

The viscoelastic dynamics of the cell membrane is described by five nonlinear partial differential equations written for five medium functions (*h* - membrane deflection along the Z-axis; *T*-temperature, and *σ*-membrane surface density; two components of the hydrodynamic velocity *V*_*i*_ in the membrane plane), dependent on time and spatial coordinates:

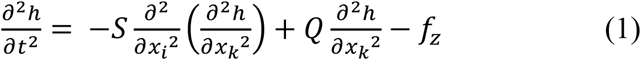

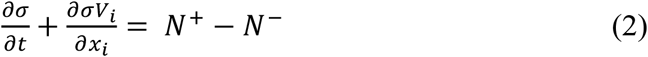

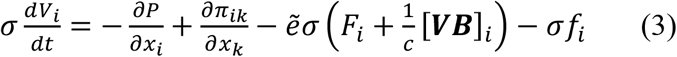

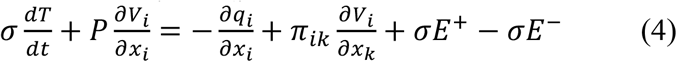

where

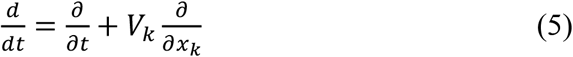

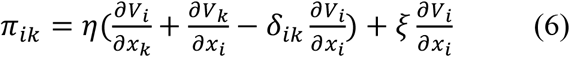

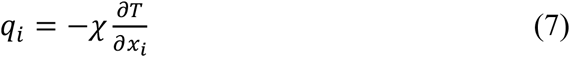

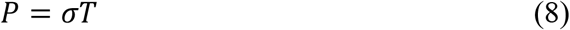

where *S* is the membrane stiffness; *Q* is its elastic tension; *f*_*z*_ is external forces perpendicular to the membrane; *F*_*i*_ is the electric field intensity;**B** is the magnetic induction vector; *f*_*i*_ are non-electromagnetic external forces along the membrane; *P* is pressure; *π*_*ik*_ is the viscosity stress tensor; *η* is the coefficient of dynamic viscosity; *ξ* is the coefficient of volume viscosity; *q*_*i*_ is the heat flow vector; χ is the thermal conductivity coefficient; *N*^*+*^, *N*^*−*^ are external sources and sinks of substances changing surface density (e.g., attachment of new molecules due to chemical reactions); *σE*^*+*^,*σE*^*−*^ - are external sources and sinks of thermal energy; 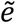 is the average charge per particle with mass *m*.

Due to the two-dimensionality of the system, indices take only two values. In the system (1-4), the electric and magnetic fields are assumed to be given by external fields. If they are variable, Maxwell’s equations must be added to system (1-4). The influence of the magnetic field on the permeability of cell membranes and other biological processes is discussed in many works (see, for example, Zablotskii et al., 2016; Barbic, 2019). Equation (3), indicating that such an influence is possible, can serve as a convenient tool for studying the effect of the magnetic field on the cell. It is important to note that the system (2-4), except for changing two coefficients from 3/2 to unity, repeats the system of equations for magnetized plasma (see, for example, Braginsky, 1963). Let’s write the system (1-4) for the Cartesian coordinate system, where the *x,y* axes are located in the membrane plane, and replace *x* _1_with *x* and *x*_*2*_ with *y*:

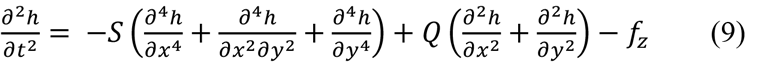

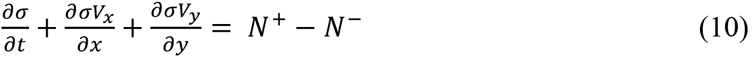

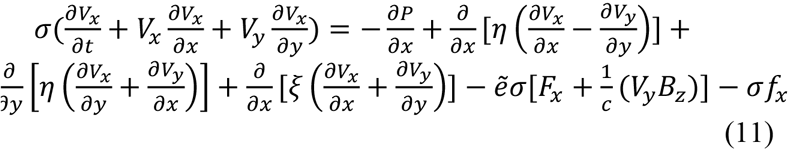

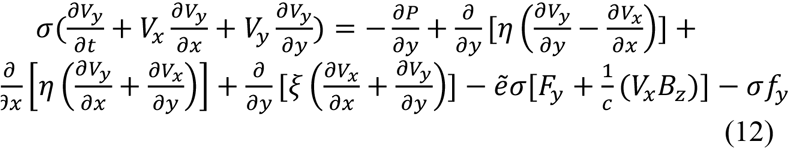

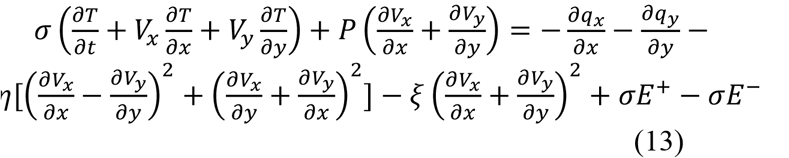

If we seek solutions to equations (9-13) as a sum of the stationary solution and small-amplitude oscillations of all quantities, for small perturbations of hydrodynamic variables, we can obtain a system of linearized differential equations that is much simpler to analyse (Fridman and Gorkavyi, 1999). Let’s write the following system of linearized equations, where external electromagnetic fields are assumed to be zero, and all quantities *h, σ,V*_*x*_,*V*_*y*_,*T,f,N,E* characterize small perturbations:

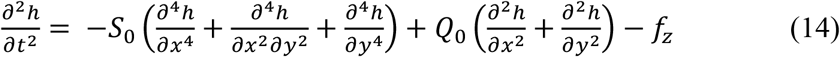

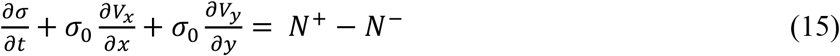

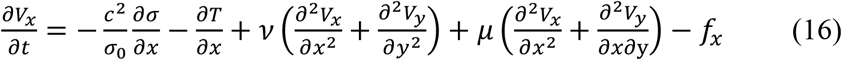

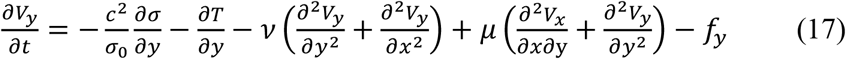

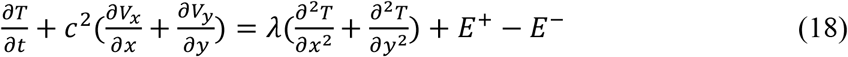

where ;*T*_0_ =*C*_2_; *λ*= *χ*_0_*/ σ*_0_;*v= η*_0_/*σ*_0_; *μ*= *ξ*_0_/*σ*_0_;– kinematic coefficients of thermal conductivity, viscosity, and volume viscosity, respectively. The zero index is assigned to the stationary quantities, and all terms in the equation consisting solely of stationary quantities have been separated into stationary balance equations (Fridman and Gorkavyi, 1999).

## 3. Dispersion Equation for Lateral Oscillations

Let’s transform equations (14-18) into a system of algebraic equations for the amplitudes of perturbations, denoted by a hat. Let’s consider only oscillations that depend on the coordinate *x*. For example, for surface density, we can write:

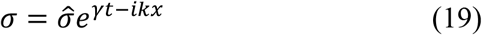

where the increment *γ*= −*iω*; and the wave vector *k* = 2*π*/*L*. If *S*_0_, *Q*_0_. *f*_*z*_ do not depend on density, temperature, and velocity, then the system (14-18) decomposes into a system of hydrodynamic equations (15-18) in the membrane plane and a separate equation (14) for transverse membrane oscillations. Since we are considering perturbations only along *x*, we can omit the symmetric equation (17) for the *y*-component of velocity. From equations (14-16, 18), let’s write a system of algebraic equations for the amplitudes of perturbations (details of obtaining such equations can be found in the book by Fridman and Gorkavyi, 1999). The main assumption of the derivation is the short-wavelength approximation: the lengths of perturbations are smaller than the characteristic spatial scale for changes in stationary quantities or the size of the cell.

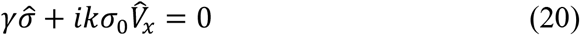

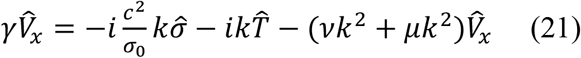

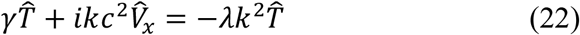

The system of equations (20-22) describes oscillations in the membrane plane for the simplest case of the absence of external forces, chemical reactions (or their analogs), and sources (sinks) of energy. Setting the determinant of the system (20-22) to zero, we obtain the dispersion equation describing the fundamental oscillations of the membrane:

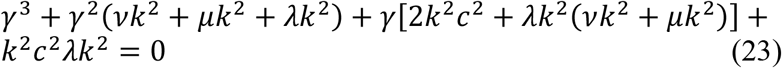

Consider individual branches of oscillations.

1. The case of sound oscillations, where *γ* ∼ *k*^2^ *c*^2^*≫ vk*^2^ + *λk*^2^. From (23), we obtain:

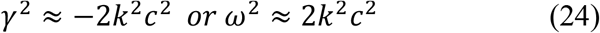

Sound oscillations in the membrane plane are stable, and the phase velocity of these oscillations *v* = ω/*k* is equal to 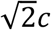.
2. The case of thermal oscillations, where *γ* ∼ λ*k*^2^ *≪ k*^2^ *c*^2^From (23), we obtain:

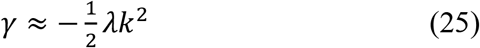

This thermal oscillation decays due to heat conduction.
3. The case of viscous oscillations:

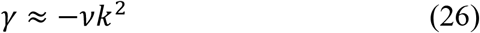

This oscillation decays due to viscous dissipation. It is worth noting that in astrophysical environments, there are cases where the diffusion coefficient is negative, and the branch of viscous oscillations may be unstable (Fridman and Gorkavyi, 1999). However, for a cellular membrane, all three considered simple branches of oscillations typically should be stable. Nevertheless, complicating the system due to the presence of external forces or sources of matter or energy alters the nature of the described branches of oscillations (often preserving their characteristic frequencies) and may lead to their instability.

If there are sources and sinks of energy in the cellular membrane that depend on the membrane density, equation (22) transforms into a more complex form:

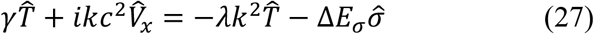

where

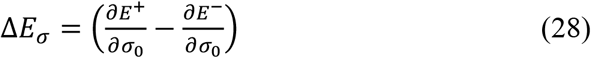

Then, in the dispersion equation (23), the last term is replaced by the following:

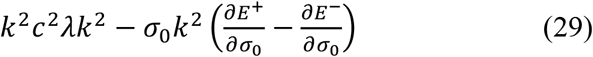

The dispersion equation for thermal oscillations takes the form:

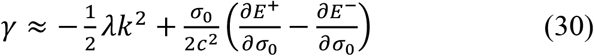

Equation (30), with a positive second term on the right-hand side, allows for instability (positive value of *γ*), representing thermal balance instability (Fridman and Gorkavyi, 1999).

Let’s write the system of algebraic equations for perturbation amplitudes in the case of chemical terms in the continuity equation. For simplification, let’s assume the isothermal case and neglect all temperature fluctuations and the energy equation:

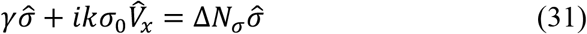

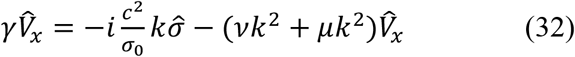

where:

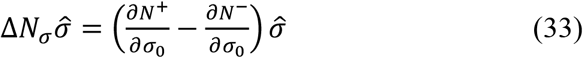

From (31)-(32), we obtain the following simple dispersion equation:

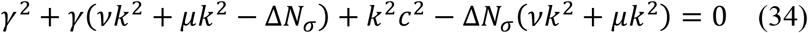

We will look for the branch of oscillations with an increment *γ*∼ Δ*N* _*σ*_. Let Δ*N* _*σ*_ ∼ *vk*^2^, and *k*^2^ *c*^2^ ≫ Δ*N* _*σ*_ (*vk*^2^ + *μk*^2^). In this case, from equation (34), a high-frequency branch of sound oscillations analogous to (24) will emerge: *γ*^2^ ≈−*k*^2^ *c*^2^. The second branch will describe chemical oscillations with diffusion:

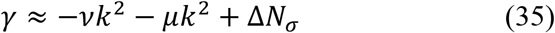

It can be unstable if the following criterion is satisfied:

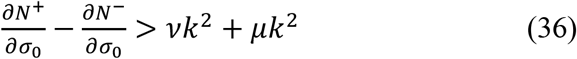

The meaning of (36) is quite simple: chemical reactions must increase the concentration of molecules in a certain zone faster than diffusion, which tends to disrupt the concentration increase.

Several instabilities can arise from corresponding terms *f*_*x*_ and *f*_*y*_ - for example, friction with the external environment or interactions of membrane lipids with embedded proteins.

If there are two substances in the cell membrane that react chemically with each other, the system of equations (1-4), written for each component, allows obtaining a system of 8 equations that will describe chemical oscillations in the diffusion system, including instabilities giving rise to spatial structures. The hydrodynamic system for each component can be reduced to a diffusion equation; such pairs of diffusion equations with chemical terms are widely discussed in the literature to describe morphogenesis processes (Murray, 1977; Belintsev, 1991) - see Fig. 1.

**Fig. 1.**
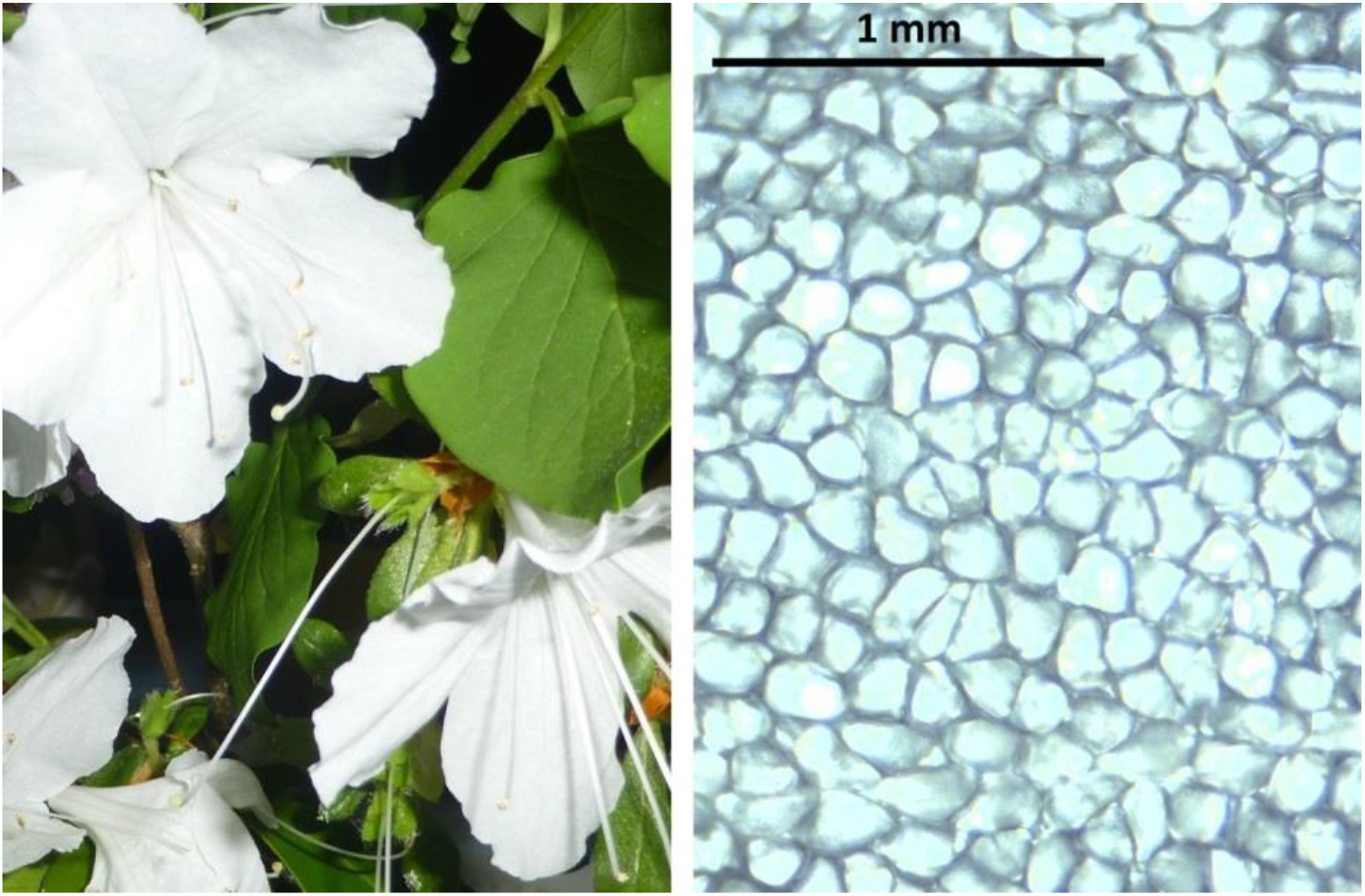
Azalea flower petal (left), under a microscope, appears as a pattern of well-distinguished cells formed as a result of morphogenesis (right). The photo on the right was obtained using an optical microscope.

## 4. Dispersion equation for bending oscillations

Several studies (Ramaswamy et al., 2000; Zhang, 2009; Gefen, 2010; Hannezo et al., 2011; Fouchard et al., 2020) have been dedicated to instabilities of cell membranes, including bending instability. Let’s consider transverse oscillations described by equation (14). In the absence of external forces, the dispersion equation for transverse oscillations of the membrane can be derived from (14) as follows:

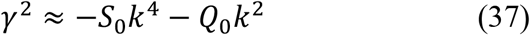

For the case of *S*_0_*k*^4^ ≫ *Q*_0_*k*^2^, we obtain from (37) the equation for the real frequency of transverse oscillations of the membrane, determined only by its stiffness:

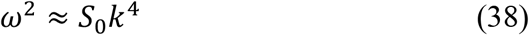

In the opposite condition *S*_0_*k*^4^ ≪ *Q*_0_*k*^2^, we obtain from (37) the equation for the frequency of transverse oscillations, determined only by the tension of the membrane:

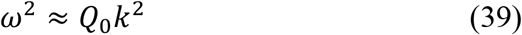

In usual conditions, *S*_0_ and *Q*_0_ are positive, thus (38) and (39) describe oscillations with a real frequency. From (39), we obtain the phase velocity of wave propagation along the membrane with dominant tension: 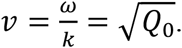. For stiff membranes, from (38), we get 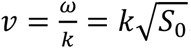. Thus, the speed of bending oscillations will depend not only on the membrane parameters but also on the wavelength: bending oscillations with shorter wavelengths will propagate faster.

The elastic tension of the membrane, *Q*, stabilizes transverse membrane oscillations — similar to how the elasticity of a rubber sheet prevents the growth of transverse waves because they stretch the sheet. However, there may be cases where *Q* can be negative. For example, during the withering of azalea petals, cell wrinkling occurs, and various types and scales of bending waves appear on their surface (see Fig. 2). Apparently, this process can be described by negative tension.

**Fig. 2.**
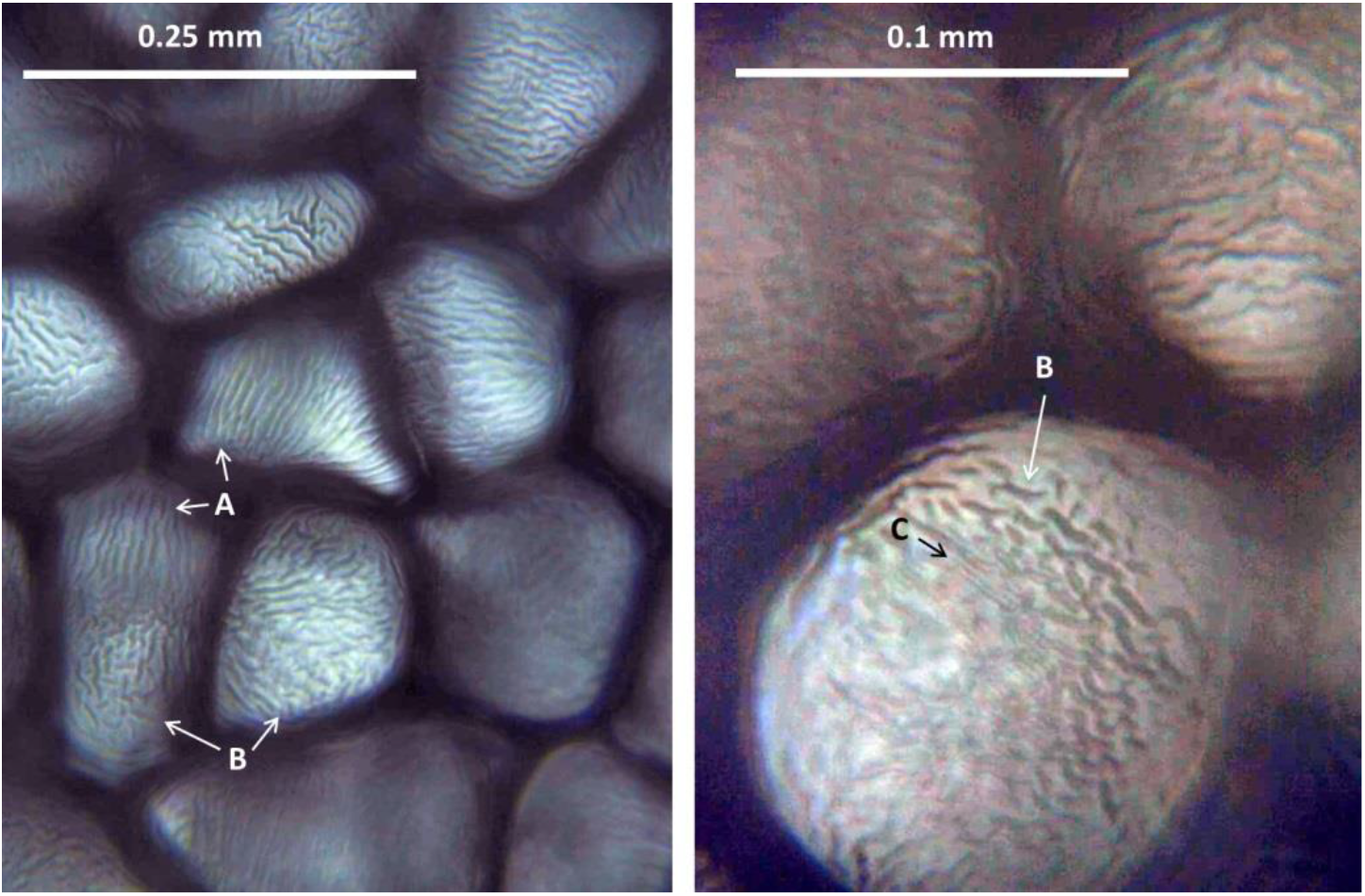
Left: cells of a withering azalea petal demonstrate a regular structure of wrinkles of various types – A and B. Right: at high magnification, different scales of wrinkles bending the cell surface are visible – B and C. Both images were acquired using an optical microscope.

The process of wave formation on the cell surface can be described by negative tension *Q*, which can induce bends in the membrane. Then, equation (37), written without considering external forces, provides the condition for such bending instability, which can only occur with negative *Q*_0_:

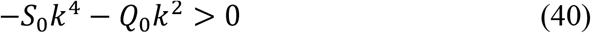

From (40), it immediately follows that the scale of the bending instability of the erythrocyte membrane will depend on the membrane stiffness as follows:

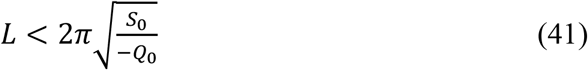

Interestingly, the criterion for bending instability (41) allows for various interpretations: it holds for both a negative parameter *Q* - the elastic tension of the membrane, and a negative *S* - its stiffness.

The egress of malaria parasites *P. falciparum* from infected erythrocytes, extensively studied by the K. Brown-Bretton group at the University of Montpellier (Abkarian et al., 2011; Chandramohanadas et al., 2011), involves the following physical processes: a. the internal membrane layer of the infected erythrocyte becomes supersaturated with substances and elastic energy, leading to the formation of irregularities on the membrane; b. after the appearance of a pore in the membrane, the edges of the pore begin to expand and bend. As a result, the cell literally turns inside out, releasing parasites into the intercellular space (Abkarian et al., 2011; Chandramohanadas et al., 2011). It is logical to expect that the general dynamic equations (1), (14), and (37) should describe such bending instability of the membrane of an infected erythrocyte. The hypothesis that such turning inside out of the erythrocyte can be characterized by negative membrane stiffness is attractive. This would maintain equation (41), were the positivity of the square root expression would be determined not by the negativity of stiffness *S*_0_.

Such a hypothesis can be experimentally tested by altering the stiffness of the cell membrane, for example, by adding cholesterol (Reigada et al., 2010; Pinkwart et al., 2019), and measuring the characteristic scales of swelling of the supersaturated membrane.

Taking external forces into account expands the range of processes described by equation (37). For instance, considering the elasticity *E*_*el*_ of the medium surrounding the membrane introduces an external force stabilizing the oscillations: *f*_*z*_ = *E*_*el*_ *h*. If the external medium moves relative to the membrane, the external force can induce the growth of bending oscillations — similar to how wind causes waves on water. If the external medium has dynamics independent of the surface film, its equations should be written separately, and their solutions considered in the film’s dynamics as boundary conditions and external perturbing forces. The most complex case arises when the influence of the film or membrane on the external medium cannot be neglected. In this case, both sets of equations need to be solved simultaneously.

Let’s consider the relevant and typical case of the influenza virus entering a cell when it has a quasi-spherical shape and induces the bending of the membrane (Chandramohanadas et al., 2011), similar to endocytosis processes. Let’s expand the external force in terms of the small parameter of deflection *h*:

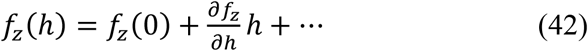

From (9) and (37), we obtain the following dispersion equation:

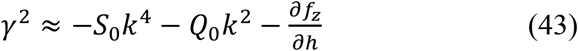

If we compare half the wavelength of the bending perturbation of the membrane *L*/2with the diameter of the quasi-spherical virus 2*R*(or the object being engulfed by the cell during endocytosis), we get an estimated condition for the membrane deflection during virus entry into the cell (assuming the positivity of 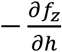):

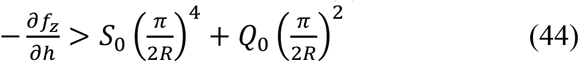

From (44), it follows that meeting this condition becomes challenging with an increase in the stiffness of the cell membrane and/or its elastic tension. Increasing these membrane characteristics should make it more resistant to the penetration of various viruses.

## 5. Conclusion

The system of differential equations (1-4), obtained through classical methods developed for transport theory, describes the dynamics of cell membranes and monomolecular films, taking into account viscosity (diffusion), pressure, thermal conductivity, energy sources, chemical reactions (or their equivalents), and any external or internal forces. Transverse deviations depend on elastic tension and stiffness of the cell membrane, as well as external forces, such as the elasticity of the surrounding medium or the influence of an external object like a virus.

The partial differential equations for a complex system are significantly simplified when considering a non-equilibrium system close to a steady state. In such a case, the system of differential equations is linearized and transformed into a system of algebraic equations, from which a dispersion equation can be derived to describe stable and unstable oscillations of cell membranes.

## Acknowledgements

This work was supported by the grant of the Region Languedoc-Roussillon for invited professors and was partially accomplished during a monthly stay of author at the University of Montpellier as a Visiting Professor.

